# A self-initiated two-alternative forced choice paradigm for head-fixed mice

**DOI:** 10.1101/073783

**Authors:** Fred Marbach, Anthony M. Zador

## Abstract

Psychophysical tasks for non-human primates have been instrumental in studying circuits underlying perceptual decision-making. To obtain greater experimental flexibility, these tasks have subsequently been adapted for use in freely moving rodents. However, advances in functional imaging and genetic targeting of neuronal populations have made it critical to develop similar tasks for head-fixed mice. Although head-fixed mice have been trained in two-alternative forced choice tasks before, these tasks were not self-initiated, making it difficult to attribute error trials to perceptual or decision errors as opposed to mere lapses in task engagement. Here, we describe a paradigm for head-fixed mice with three lick spouts, analogous to the well-established 3-port paradigm for freely moving rodents. Mice readily learned to initiate trials on the center spout and performed around 200 self-initiated trials per session, reaching good psychometric performance within two weeks of training. We expect this paradigm will be useful to study the role of defined neural populations in sensory processing and decision-making.

## 1. Introduction

A central goal of neuroscience is to identify the neural circuits underlying cognitive behavior. Pioneering research has been done recording from macaque monkeys engaged in psychophysical two alternative choice (2AFC) tasks. These recordings made it possible to directly correlate cognitive variables with neural activity, thereby identifying candidate regions for sensory integration, decision making and movement planning (Parker and Newsome 1998; Romo and Salinas 2001; Gold and Shadlen 2007). However, identifying the neural circuits engaged in these tasks has been held back by the large experimental overhead in primate research.

In order to increase experimental flexibility, we and others have developed 2AFC sensory discrimination tasks for freely moving rats that were directly inspired by tasks developed for primates (Uchida and Mainen 2003; Kepecs et al. 2008; Otazu et al. 2009; Erlich et al. 2011). Importantly, these task designs preserve essential features of the original paradigms for macaques: trials are actively initiated with a ‘fixation’ action, and the subject is forced to choose one of two equal but opposite actions on every trial to indicate its decision. These features ensure a consistent internal state at the onset of each trial and decouple the subject’s choice from its motivational state, allowing us to attribute error trials to perceptual or decision errors, as opposed to changes in task engagement.

There are now hundreds of transgenic mice available that grant access to specific cell populations (Gerfen et al. 2013; Taniguchi et al. 2011), providing increasing incentive to develop psychophysical tasks for mice. We have recently shown that freely moving mice can reach similar performance as rats in an adaptive decision-making task (Jaramillo and Zador 2014). However, in order to fully leverage the transgenic and viral toolkit, several groups have developed head-fixed paradigms (Sanders and Kepecs 2012; Mayrhofer et al. 2013; Harvey et al. 2012; Guo et al. 2014b; Resulaj and Rinberg 2015; Burgess et al. 2016) that allow two-photon calcium imaging and optogenetic manipulations of specific cell populations during behavior. However, it has been difficult to include active trial initiation in these paradigms to disambiguate perceptual errors from mere lapses in attention.

**Figure 1:**
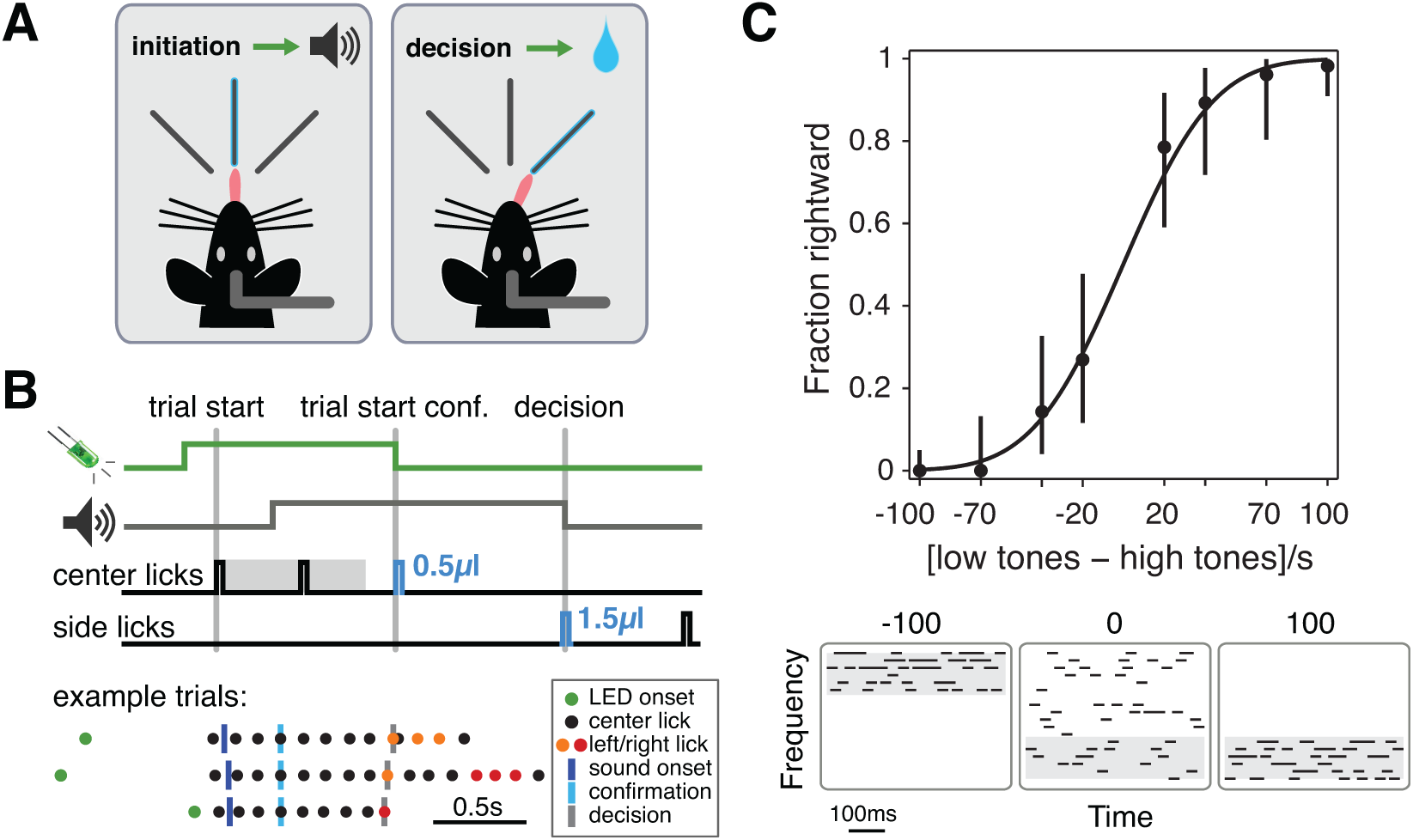
‘3-lick’ 2AFC paradigm for head-fixed mice. **A**: Schematic of the setup (seen from above) **B**: *(top)* Schematic of trial structure. After LED onset, mice could initiate a trial by licking the center spout, triggering the stimulus (with a delay). Mice were required to lick until the end of ‘fixation’ period (grey shading), which triggered a small reward and turned off the LED. The first side lick constituted the response and turned off the stimulus. *(bottom)* Three example trials with recorded task events (see legend). Trials were aligned to the trial start confirmation lick on the center spout (light blue). **C:** Best single session psychometric curve from mouse fu032. Data points show mean with 95% confidence intervals, fit with a cumulative Gaussian distribution (see Methods/Data analysis). Example 0.5s long stimuli for high frequency trials are plotted below, with the target octave shaded gray (stimulus -100: 20kHz-40kHz; stimulus 100: 5kHz-10kHz; frequency axis is logarithmic).

Here we describe the ‘3-lick’ two-alternative forced choice paradigm for head-fixed mice, in which we replaced the three ports typical for freely moving paradigms with three lick spouts. In this paradigm, the mouse initiates the trial by licking the center spout for a prescribed period, after which it is free to signal its decision by licking either the left or right spout. The period of center licking is analogous to the visual fixation period used in many monkey tasks. We trained mice on a ‘cloud-of-tones’ frequency discrimination task initially designed for rats (Znamenskiy and Zador 2013; Xiong et al. 2015). Mice readily learned to initiate trials by licking the center spout, and to lick the side spouts to collect water rewards. Performance was above 90% on easy trials within two weeks of training, and varied smoothly for intermediate difficulties. The ‘3-lick’ paradigm extends existing head-fixed mouse behavior to offer active trial initiation, matching the well established 3-port paradigm for freely moving rats and mice.

## 2. Results

### ‘3-lick’ paradigm

Our goal was to train mice to perform a head-fixed, two-alternative forced-choice (2AFC) auditory discrimination task previously designed for freely moving rats (Znamenskiy and Zador 2013; Xiong et al. 2015). To this end, we modified an existing paradigm with two lick spouts for left/right reports (Mayrhofer et al. 2013; Guo et al. 2014b) to include a third, middle spout for a ‘fixation-like’ trial initiation (Fig. 1). This ‘3-lick’ paradigm thus offered two important advantages of the widely used 3-port paradigm for freely moving rodents (Uchida and Mainen 2003; Otazu et al. 2009; Erlich et al. 2011; Clark et al. 2011; Raposo et al. 2012): a 2AFC task structure, and a self-initiated trial structure with a fixation-like period during stimulus presentation. As stimulus, we used a ‘tone cloud’ as previously described (Fig. 1c, Znamenskiy and Zador (2013)).

The trial structure is depicted in Fig. 1a-b (see Methods/Task design). After the inter-trial interval, an LED signaled trial onset: licking the center spout triggered stimulus presentation and a ‘fixation’ time window. The first center lick after that window confirmed trial-initiation, signaled by a small center water reward (0.5*µ*l) and LED offset. The first side lick turned off the stimulus, and triggered a 1.5*µ*l reward if correct (see also Supplemental movie 1).

### Mice learned the task in under 2 weeks

The data presented here was collected from four mice (named fu029-fu032) trained for 21-25 sessions over 12-14 days. All four mice readily learned the basic trial structure (Fig. 2a, see Methods for detailed training procedure). First, they learned to collect free rewards on the center spout upon LED onset (>200 trials cumulative across sessions; achieved in 2-4 sessions). The latency to lick after LED onset (trial initiation) decreased over training, but remained variable from trial to trial (Fig. 2b). Mice also learned to persist licking the center spout (‘fixating’) for 300ms for a small reward (Fig. 2c).

Training mice to lick the side spouts after trial initiation posed a challenge. It took the mice 7-12 sessions to reach >200 cumulative trials where they collected free reward on the side spout after initiating a trial (Fig. 2a). All four mice performed the task at >80% correct on the easiest trials (100 tones/s target octave) on two consecutive sessions after 11-22 sessions, and three out of four mice eventually performed at >90% correct on the easiest trials on two subsequent sessions (achieved in 17-21 sessions, Fig. 2a, e).

We did not train animals long enough to achieve asymptotic performance, so as expected performance across sessions was variable. Nevertheless, performance remained mostly above 80% correct once the mice were performing the full task (Fig. 2e, only sessions with >50% full task trials shown). After learning, mice performed 258±73 trials per session, with 7±4% passive trials (no licking for 1min) and 12±7% early side lick trials (Fig. 2d).

### Mice generalized well to intermediate stimuli

Once the mice were performing above 80% on easy trials (fu029 failed to reach this stage within this time frame), we gradually introduced intermediate stimuli at three fixed stimulus difficulties (denoted as target/non-target tones/s): 85/15, 70/30 and 60/40, corresponding to ±70, ±40 or ±20 tones/s in favor of a rightward choice. We presented at most 80% intermediate trials in a given session. Performance tracked the stimulus difficulties smoothly, remaining near 90% correct on easy trials (Fig. 2f). Thus, our mice produced psychometric curves on single sessions that were qualitatively similar to those previously obtained in freely moving rats performing the tone cloud task (Znamenskiy and Zador 2013).

## 3. Discussion

We present here the ‘3-lick’ paradigm for training head-fixed mice in sensory discrimination tasks. By adding a middle spout to the existing head-fixed licking approach (Mayrhofer et al. 2013; Guo et al. 2014b), we introduced active trial initiation and a fixation period to the head-fixed 2AFC lick left/right paradigm. Mice learned to perform the task well, achieving approximately 90% correct performance on the easiest stimuli within a two-week training period. Three out of four mice yielded single session psychometric curves qualitatively similar to ones obtained from freely moving rats performing the same tone cloud task (Znamenskiy and Zador 2013). We anticipate future optimization in task structure and training procedure to further improve training success rate and learning speed.

We chose to train animals with the lick left/right approach introduced by Mayrhofer et al. (2013), as opposed to the trackball approach from Sanders et al. (2012), because in our hands mice learned more readily to report their choices by licking than by turning a ball or wheel with their front paws to the left/right (data not shown). It is not clear why mice can readily learn to control a wheel or treadmill in the context of other tasks (Burgess et al. 2016); a key feature may be the availability of real-time visual feedback about the position of the wheel (but see Resulaj and Rinberg 2015). For the cloud-of-tones and other tasks in which there is no natural way to provide such feedback, the lick left/right paradigm may be easier to train.

**Figure 2:**
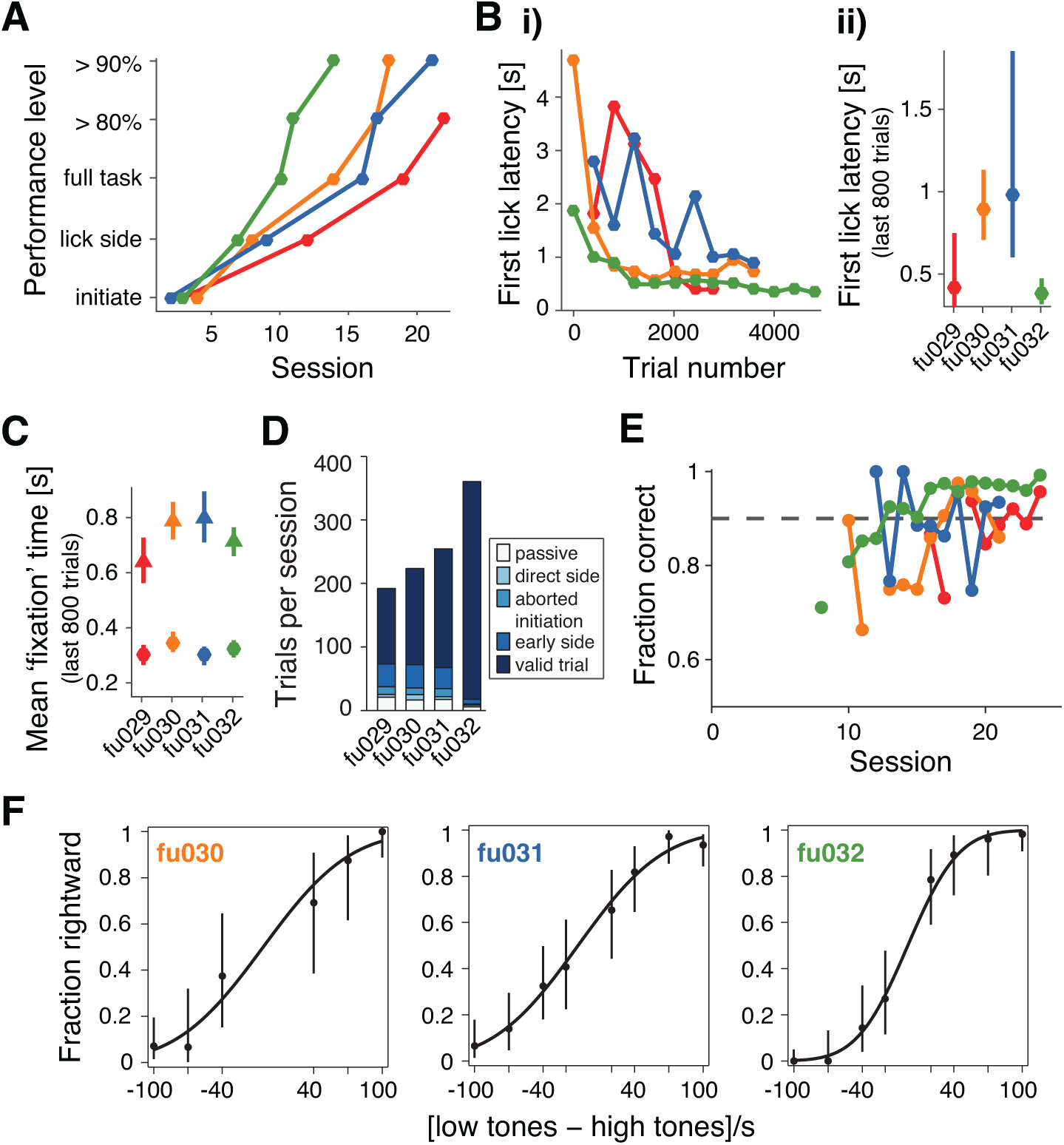
Mice reached high performance levels within 2 weeks of training. **A**: Learning across sessions as quantified with post hoc criteria. *Initiate:* >200 trials were initiated (cumulative across sessions); *lick sides:* >200 trials where mouse collected side water reward after initiation (cumulative across sessions); *full task:* >200 trials where mouse made a choice; *>80% / >90%*: the second of two consecutive sessions with average performance on easiest trials >80% / >90%. **B**: i) Latency to lick after LED onset (400 trials bins). ii) Median and 0.25/0.75 quartiles for last 800 trials. **C**: Mean and standard deviation of time from first center lick to center reward (circles, trial start confirmation) and first center lick to first side lick (triangles, valid trials only). Triangles also approximate sound duration (absent the small delay from trial start to sound onset). **D**: Mean number of trials per session (last 6 sessions). *passive:* no licks for 1min; *direct side:* side lick before center lick; *aborted initiation:* did not confirm trial start; *early side:* side lick before trial start confirmation; *valid trial:* initiated trial correctly and made a choice. **E**: Performance on easiest trials across sessions. Only sessions with >50% full task trials are shown, and only valid trials are taken into account. **F**: Best single session psychometric curves for each mouse (fu029 did not reach this stage). Mean and 95% confidence intervals are shown, together with cumulative Gaussian fit. fu030: 193 trials; fu031: 342 trials; fu032: 343 trials.

The addition of the central spout brings two improvements with it over existing lick left/right paradigms: first, animals can actively initiate trials, and second, through the use of a small reward they can readily be trained to ‘fixate’ during trial initiation by persistent licking on the center spout. Active trial initiation is important because it acts to reduce the number of trials in which the animal makes a wrong choice due to lack of task engagement. Whenever the animal initiates, it is likely that it is motivated to perform the next trial, so that errors are more likely to be perceptual or decision errors than mere lapses in attention. Indeed, our mice initiated trials with variable latency (Fig. 2b), and were sometimes passive for prolonged periods before doing more trials (Fig. 2d, passive trials).

Both in the freely moving and head-fixed paradigm, the fixation period is key to avoid impulsive behavior that leads to hasty, uninformed choices. However, the fixation period can be hard to enforce (but see Guo et al. 2014a; Burgess et al. 2016). Delivering a small reward at the end of the fixation period facilitates this training process. Here we required only that mice fixate for 300ms, but it is likely that this period could readily be extended to seconds. This could be of interest for working memory tasks in which stimuli are presented sequentially (separated by seconds) to be compared by the subject.

There are still several aspects of the current task design that fall short of the freely moving paradigm. Most notably, our mice only performed around 200-300 trials per session (Fig. 2d), around three fold fewer than freely moving mice and rats (Jaramillo and Zador 2014). One option to increase trials per session is to replace water with sucrose solution (Guo et al. 2014a), food deprivation (soy milk as reward, (soymilk as reward, Poort et al. 2015) or optogenetic stimulation of the VTA (Burgess et al. 2016). Another shortcoming is the lick detection in its current form, which relies on grounding the mouse and is therefore unsuitable for electrical recordings; piezo-based circuits solve this problem (Mayrhofer et al. 2013). It is also worth noting that the stimulus presentation occurs while the animal is repeatedly licking the center spout (Fig. 1b). However, we believe the benefit of trial initiation and training efficiency will usually outweigh concerns such as interaction of sensory processing and licking, or licking-induced motion artifacts. Lastly, the training procedure is more cumbersome than in the freely moving paradigm (see Methods/Training procedure). The main issue in our hands is the lack of exploratory side licks in the first few days of training. Mayrhofer et al. (2013) solved this problem by allowing for small head rotations during initial training so as to make the side licking less challenging. We envisage a similar approach, consisting in moving the side lick ports laterally with a motor to initially decrease the distance to reach the side spouts.

In summary, the ‘3-lick’ paradigm is a useful addition to existing head-fixed mouse psychophysics paradigms, in particular because of its active trial initiation and fixation period. In addition, the center spout may provide a good approach to efficiently train head-fixed mice on working memory paradigms with long waiting periods.

## 4. Methods

### Animal subjects

Animal procedures were approved by the Cold Spring Harbor Laboratory Animal Care and Use Committee and carried out in accordance with National Institutes of Health standards. Four 8-week-old male CBA/CaJ wild type mice (The Jackson Laboratory, stock #000654) were used for training. Mice had free access to food, but were on a water restriction schedule of 1ml per day. Weight was monitored daily, and animals that exhibited weight drop below 80% of their initial weight were supplemented with additional water. For head-fixation, mice were implanted with a custom designed lightweight titanium head bar. They were anesthetized with ketamine/medetodomine (60/0.5 mg/kg). An additional analgesic was injected (meloxicam 2mg/kg) and the mice were placed in a stereotaxic apparatus. The scalp was removed above the entire cortical area, the skull cleaned with hydrogen peroxide and then covered in several layers of metabond adhesive (parkell, S380) and let to dry for 10min. The head bar was attached to the metabond with dental acrylic (Lang, Jet denture repair powder/liquid). Mice were allowed to recover for several days before starting the water restriction schedule. Mice were head-fixed using custom designed clamps positioned over a vertical wheel (3D printed) running on ball bearings held by Thorlabs equipment (Thorlabs, New Jersey). For details please consult parts description and parts list in the Supplement.

### Behavioral apparatus

For details please consult parts description and parts list in the Supplement. Training was done inside custom sound-booths by Industrial Acoustics Company (Bronx, New York). Water was delivered through 19 gauge stainless steel tubing connected via rubber tubing (Silastic) to solenoid valves (Lee company) located outside the sound box. Water volume was calibrated to 0.5*µ*l on the center spout and 1.5*µ*l on the side spouts, corresponding to valve times of 2ms and 6ms respectively. The three spouts were held in a 3D printed holder, each connected to a custom lick detection circuit. Mice were grounded over their head-fixation gear, closing the respective circuit whenever they licked one of the spouts, thereby triggering reward delivery or other behavioral events. Sound was calibrated and delivered as described previously (Jaramillo and Zador 2014). The behavioral system was controlled by a custom Matlab (Mathworks) program running within the BControl framework (http://brodylab.princeton.edu/bcontrol; see also https://sanworks.io for behavior software currently in use).

### Task design

The trial schematic is sketched in Fig. 1b, with three actual example trials below. Mice were trained to withhold licking during the inter-trial interval (ITI, drawn from an exponential distribution of 2s mean, truncated at 3s, then offset by a constant 1.5s; notation: [1.5/2/3]). Any ITI licks restarted the ITI. After the ITI, an LED signaled trial onset and remained on for at most 1min or until the mouse successfully initiated a trial. The first lick on the center spout triggered both presentation of the tone cloud stimulus ([0.05/0.05/0.05]; see above for notation), and a fixation-like time window (fixation time, [0.2/0.15/0.3]) during which mice were not allowed to lick the side spouts yet. The first center lick after fixation time confirmed trial-initiation, triggering a small water reward (0.5*µ*l) at the center spout and turning off the LED. Mice were now allowed to make a choice (within 2s) by licking either of the side spouts. Any side licks before this point aborted the trial, jumping to the ITI with an added early lick timeout (3s). Correct choices were rewarded with a 1.5*µ*l water reward, errors punished with a mild air puff to the snout and an error timeout (4s). The sound was turned off at the first side lick.

### Stimulus design

We used the same ‘tone cloud’ stimulus as described recently (Fig. 1c, Znamenskiy and Zador (2013)). The tone cloud consisted of a series of 30ms long pure tones presented at a fixed rate of 100 tones per second. The frequencies of these tones were drawn from 18 logarithmically spaced slots spanning three octaves (5kHz to 40kHz). Each tone was ramped up/down for 3ms to avoid spectral splutter during playback. On easy trials, frequencies were drawn exclusively from the target octave: on rightward trials from the low octave (5kHz-10kHz), and on leftward trials from the high octave (20kHz-40kHz). On intermediate trials, some of the tones were drawn from the two non-target octaves. For example, a stimulus strength of 80 meant that for each time slot, there was a 80% chance for the tone to be picked from the target octave, and a 20% chance for it to be picked from the other two octaves. Sound intensity of individual tones was kept constant on a given trial, but was drawn from a uniform distribution from 50-70dBSPL across trials to discourage mice from using loudness for discrimination.

### Training procedure

#### First stage: initiate trials on center spout

We presented mice with simple trials: ITI, then LED onset together with free water on the center. A trial was correct if the mouse licked off the water within 2s. As soon as mice were engaged, we dispensed water only upon contact and started delaying the delivery (increasing fixation time), forcing mice to persist licking the center. Technically, mice could have licked once the center, waited for the fixation time to elapse, then licked once more the center to confirm trial initiation; in practice, they licked at a fixed rate of roughly 8Hz throughout fixation time.

#### Second stage: lick side spouts after initiation

This resembled the full task, but water was dispensed automatically from the correct side upon trial initiation. Mice were initially shown manually with a pipette tip that water was available on the side spout, at the beginning of each session over several trials. As they started exploring more, the probability of free water trials (‘direct trials’) was gradually reduced, and trials were introduced where mice still got reward if they first licked the wrong side and then corrected their choice (‘next correct trials’). Direct and next correct trials were then phased out and mice had to lick only the correct side to get a reward; i.e. they were doing the full task.

#### Third stage: psychometric curve

Once a given mouse was performing >80% correct, we gradually introduced psychometric trials (intermediate difficulties).

#### Remarks

On most days, mice were trained in two sessions of around 30min each, as they frequently stopped working at <0.5ml total reward in a given session. A session always started with a few easy trials with direct water delivery on the correct side to engage the mouse. Throughout training, premature side licks and licks during the ITI aborted the trial. Mice tended to exhibit strong side biases before they were experts at the task. We did three things to keep these in check: limit the number of consecutive trials on one side initially to 2, and increase the number of trials and the reward size on the mouse’s weak side. While mice were not yet performing steadily, we increased the reward size at most three-fold (4.5*µ*l).

### Data analysis

Analysis was done in Matlab (Mathworks) with custom written routines.

Fig. 2a: We applied post hoc determined criteria as follows: initiate – >200 trials were initiated (cumulative across sessions); lick sides – >200 trials where mouse collected side water reward after initiation (cumulative across sessions); full task – >200 trials where mouse made a choice, i.e. no direct or next correct water delivery or intervention by experimenter (see above); >80% / >90% – the second of two consecutive sessions with average performance on easiest trials >80% / >90%.

Fig. 2d: Trial counts were averaged over the last 6 session of each mouse. Description of categories: passive: no licks for 1min; direct side: licked the side before the center; aborted initiation: started licking the center but did not persist post fixation time; early side: started licking the center but licked side before end of fixation time; valid trial: initiated trial correctly and made a correct/incorrect side lick.

Fig. 2f: Performance from sessions with intermediate stimuli were fit using a generalized linear model with a binomial distribution function and a *probit* link function, using the function *glmfit* in Matlab (yielding a bias and slope term per fit). Example sessions were ‘cherry picked’ for each mouse. For each stimulus difficulty, 95% confidence intervals are displayed (obtained with the function *binofit* in Matlab).

## 5. Acknowledgments

We would like to thank Uri Livneh and Priyanka Gupta for their insights and help for designing this paradigm. Josh Sanders provided code and support during development. Santiago Jaramillo provided code and tips for training. Petr Znamenskiy designed the initial tone-cloud task. Barry Burbach helped get/keep things running smoothly. Rob Eifert is our machining expert. Josh Cohen helped with training.

### 6. Conflict of interest

The authors declare no conflict of interests.

### 7. Supplemental material

The Supplemental material contains

- SupplementalMovie1.mov
- SupplementalMovie1Description.pdf
- 3lickSetup.pdf
- 3lickPartsList.pdf
- 3Dprint_files.zip

## References

C. P. Burgess, N. Steinmetz, A. Lak, Z.-H. Peter, A. Ranson, M. Wells, S. Schroeder, E. A. K. Jacobs, C. Bai Reddy, S. Soares, J. F. Linden, J. J. Paton, K. D. Harris, and M. Carandini. High-yield methods for accurate two-alternative visual psychophysics in head-fixed mice. bioRxiv, 2016. doi: 10.1101/051912.

R. E. Clark, P. Reinagel, N. J. Broadbent, E. D. Flister, and L. R. Squire. Intact Performance on Feature-Ambiguous Discriminations in Rats with Lesions of the Perirhinal Cortex. Neuron, 70(1): 132–140, 2011. ISSN 08966273. doi: 10.1016/j.neuron.2011.03.007.

J. C. Erlich, M. Bialek, and C. D. Brody. A cortical substrate for memory-guided orienting in the rat. Neuron, 72(2):330–43, 10 2011. ISSN 1097–4199. doi: 10.1016/j.neuron.2011.07.010.

C. R. Gerfen, R. Paletzki, and N. Heintz. GENSAT BAC cre-recombinase driver lines to study the functional organization of cerebral cortical and basal ganglia circuits. Neuron, 80(6):1368–1383, 2013. ISSN 10974199. doi: 10.1016/j.neuron.2013.10.016.

J. I. Gold and M. N. Shadlen. The neural basis of decision making. Annual review of neuroscience, 30:535–74, 1 2007. ISSN 0147–006X. doi: 10.1146/annurev.neuro.29.051605.113038.

Z. V. Guo, S. A. Hires, N. Li, D. H. O’Connor, T. Komiyama, E. Ophir, D. Huber, C. Bonardi, K. Morandell, D. Gutnisky, S. Peron, N.-l. Xu, J. Cox, and K. Svoboda. Procedures for behavioral experiments in head-fixed mice. PloS one, 9(2):e88678, 1 2014a. ISSN 1932–6203. doi: 10.1371/journal.pone.0088678.

Z. V. Guo, N. Li, D. Huber, E. Ophir, D. Gutnisky, J. T. Ting, G. Feng, and K. Svoboda. Flow of cortical activity underlying a tactile decision in mice. Neuron, 81(1):179–94, 1 2014b. ISSN 1097–4199. doi: 10.1016/j.neuron.2013.10.020.

C. D. Harvey, P. Coen, and D. W. Tank. Choice-specific sequences in parietal cortex during a virtual-navigation decision task. Nature, 484(7392):62–8, 4 2012. ISSN 1476–4687. doi: 10.1038/nature10918.

S. Jaramillo and A. M. Zador. Mice and rats achieve similar levels of performance in an adaptive decision-making task. Frontiers in Systems Neuroscience, 8(September):1–11, 9 2014. ISSN 1662–5137. doi: 10.3389/fnsys.2014.00173.

A. Kepecs, N. Uchida, H. a. Zariwala, and Z. F. Mainen. Neural correlates, computation and behavioural impact of decision confidence. Nature, 455(7210):227–231, 2008. ISSN 0028–0836. doi: 10.1038/nature07200.

J. M. Mayrhofer, V. Skreb, W. von der Behrens, S. Musall, B. Weber, and F. Haiss. Novel two-alternative forced choice paradigm for bilateral vibrotactile whisker frequency discrimination in head-fixed mice and rats. Journal of neurophysiology, 109(1):273–84, 1 2013. ISSN 1522–1598. doi: 10.1152/jn.00488.2012.

G. H. Otazu, L.-H. Tai, Y. Yang, and A. M. Zador. Engaging in an auditory task suppresses responses in auditory cortex. Nature neuroscience, 12(5):646–54, 5 2009. ISSN 1546–1726. doi: 10.1038/nn.2306.

A. J. Parker and W. T. Newsome. SENSE AND THE SINGLE NEURON: Probing the Physiology of Perception. Annual review of neuroscience, 1998.

J. Poort, A. G. Khan, M. Pachitariu, A. Nemri, I. Orsolic, J. Krupic, M. Bauza, M. Sahani, G. B. Keller, T. D. Mrsic-Flogel, and S. B. Hofer. Learning Enhances Sensory and Multiple Non-sensory Representations in Primary Visual Cortex. Neuron, 86(6):1478–1490, 6 2015. ISSN 08966273. doi: 10.1016/j.neuron.2015.05.037.

D. Raposo, J. P. Sheppard, P. R. Schrater, and A. K. Churchland. Multisensory decision-making in rats and humans. The Journal of neuroscience: the official journal of the Society for Neuroscience, 32 (11):3726–35, 3 2012. ISSN 1529-2401. doi: 10.1523/JNEUROSCI.4998-11.2012.

A. Resulaj and D. Rinberg. Novel Behavioral Paradigm Reveals Lower Temporal Limits on Mouse Olfactory Decisions. Journal of Neuroscience, 35(33):11667–11673, 2015. ISSN 0270–6474. doi: 10.1523/JNEUROSCI.4693-14.2015.

R. Romo and E. Salinas. Touch and go: decision-making mechanisms in somatosensation. Annual review of neuroscience, 24:107–37, 2001. ISSN 0147–006X. doi: 10.1146/annurev.neuro.24.1.107.

J. Sanders and A. Kepecs. Choice Ball: a response interface for psychometric discrimination in head-fixed mice. Journal of neurophysiology, (September 2012):3416–3423, 9 2012. ISSN 1522–1598. doi: 10.1152/jn.00669.2012.

H. Taniguchi, M. He, P. Wu, S. Kim, R. Paik, K. Sugino, D. Kvitsani, Y. Fu, J. Lu, Y. Lin, G. Miyoshi, Y. Shima, G. Fishell, S. B. Nelson, and Z. J. Huang. A Resource of Cre Driver Lines for Genetic Targeting of GABAergic Neurons in Cerebral Cortex. Neuron, 71 (6):995–1013, 2011. ISSN 08966273. doi: 10.1016/j.neuron.2011.07.026.

N. Uchida and Z. F. Mainen. Speed and accuracy of olfactory discrimination in the rat. Nature neuroscience, 6(11):1224–9, 11 2003. ISSN 1097–6256. doi: 10.1038/nn1142.

Q. Xiong, P. Znamenskiy, and A. M. Zador. Selective corticostriatal plasticity during acquisition of an auditory discrimination task. Nature, 3 2015. ISSN 1476–4687. doi: 10.1038/nature14225.

P. Znamenskiy and A. M. Zador. Corticostriatal neurons in auditory cortex drive decisions during auditory discrimination. Nature, 497 (7450):482–5, 5 2013. ISSN 1476–4687. doi: 10.1038/nature12077.

